# MirAge, a novel Matrix Analyzer for fast DNA classification

**DOI:** 10.64898/2025.12.02.691787

**Authors:** M Pater, M D de Jong

## Abstract

**Summary:** To understand the composition of metagenomics samples, fast computer tools analyze large datasets to compose a taxonomic overview. With large datasets, it is still a challenge to achieve high precision and sensitive results in fast manner. Here we show the new tool MirAge which implements a newly developed classification strategy for analyzing DNA and RNA samples. MirAge is a highly precise and sensitive tool which can be easily used to create results from large amount of data in a fast manner. The ability to combine different databases, run on complex High Performance Computing (HPC) environments and customize many settings shows MirAge to be versatile.

**Availability:** The software is freely available at https://github.com/mirdesign/mirage

## 1 Introduction

Metagenomic sequencing is an important method to study host associated and environmental microbial communities (I Sharon *et al*., 2013). Using analysis on the extracted genetic material enables taxonomic binning of sequencing reads, which is an important step in understanding the composition and function of microbial communities (Sczyrba, A *et al*., 2017).

Here, we present a new algorithm for rapidly assigning reads to reference sequences in large common used or custom DNA datasets. This novel standalone identification algorithm pursues an approach without domain specific interpretation of the data and utilizing a dot matrix comparison to calculate scores. Its implementation in the “MirAge” tool proved to be a fast and accurate solution when compared to existing methods for metagenomics read binning, representing the primary defined pillars usability, fast operation and fresh approach.

## 2 Methods

MirAge determines the taxonomic assignment of each query sequence by mapping them to a reference sequence database. Each combination of query and reference sequence (and its reverse complement) is scored using a dot matrix comparison. Each dot represents a nucleotide match. Per diagonal of the matrix, we catalog each contiguous region of dots and note the region length (*L_R_*) if it exceeds a preset threshold of *T*.

The sum of all triangular numbers 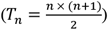 per region is divided by the sum of all *L*_*R*_. A penalty is given to nucleotides of the query when they do not match with the reference within any used region. Therefore, the number of matched nucleotides (*N*_*m*_) is divided by the length of the query sequence (*L*_*s*_).

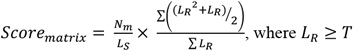

A score obtained from a region equal to *L*_*s*_ represents the maximum obtainable score (*score*_*max*_). The ratio of *score*_*max*_ out of *score*_*matrix*_, both converted with natural logarithmic to pronounce lower scored sequences more, is the normalized representation *score_final_*. This normalization provides mutual comparability of MirAge scores, where closer to 1 means a higher probability of the query originating from the designated reference.

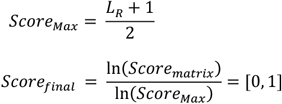

The suggested threshold for MirAge is the minimal amount of consecutive matching nucleotides which are considered to be non-random. Lowering the threshold can lead to higher number of false assignments, while increasing can result in missing assignments. Therefore, based on experience, a minimum of 20 is advised and a threshold of 31 represents a good balance between results and performance.

### 2.1 Validation

To validate the analyzing capabilities, we compared MirAge to existing tools using the CAMI 1 datasets (Sczyrba, A *et al*., 2017) and well known and described simulated datasets to benchmark accuracy and speed. The 8 simulated datasets are 7 samples from the CLARK-S manuscript (Ounit *et al*., 2016) and one Illumina HiSeq dataset used to benchmark Kraken (Wood *et al*., 2014). The RefSeq database (O’Leary *et al*., 2016) was downloaded from NCBI (release version 200;74 GB with 18049 species and 35843 genome sequences). CLARK (Ounit *et al*., 2015) and Kraken2 (Wood *et al*., 2019) were used as comparison tools as they perform equal type of analysis and state to provide speed, high precision and sensitivity. With Kraken2, the provided reference database was used. For both MirAge and CLARK, we built an index database with the NCBI reference database.

MirAge used a threshold of 31, conform to the 31 k-mer size used by CLARK.

## 3 Implementation

The algorithm has been implemented in a multithreaded C++ program. Compilation on both Windows and Unix environment is supported without code adjustments and provides a choice between a low memory or high speed mode. Using the application is straightforward and highly configurable to give expert users a high degree of customizability. The application is capable of monitoring multiple instances of itself using only a shared filesystem to communicate, guaranteeing easy deployment in HPC environments.

A separate database index is built with the sequences in the reference database, to enable fast searches. MirAge automatically applies pre-filters to only load minimal information necessary for specific analyses to further increase performance. Multiple databases can be loaded simultaneously, combining specialized databases (i.e. antibiotic resistance sequences, an *Escherichia coli* only database or some specific viral databases).

The program can run in a fast or the default sensitive mode. In fast mode, even more information is filtered out by the pre-filter, where index search results are used to limit data retrieval from disk and also decrease the number of calculations.

## 4 Results

All tools were set to use the default mode. The first analysis used 7 simulated datasets to compare CLARK and MirAge. The results showed MirAge to be much more sensitive and MirAge achieved a higher precision on 6 out of 7 samples (Figure 1D).

**Figure 1.**
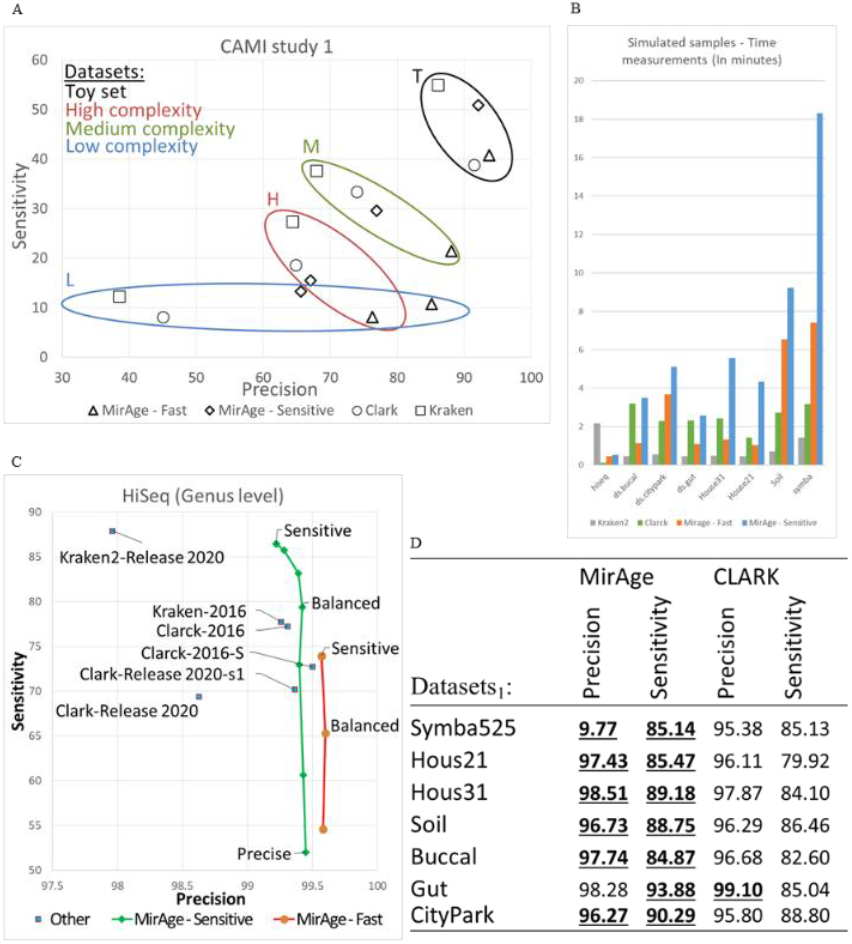
**A)** Results of analyzing CAMI 1 study samples **B)** Results of performance tests **C)** Reported and rerun results of analyzing 1 simulated HiSeq sample from Kraken **D)** Analysis results of 7 simulated samples (higher is underlined). ^1^Dataset samples from (Ounit *et al*., 2016).

The CAMI samples were used to compare CLARK, Kraken2 and MirAge (see Figure 1A). For the low complexity CAMI sample, MirAge achieved a very high precision and comparable sensitivity as the other tools. For an even higher precision, MirAge can be run in fast mode. With the medium and high samples, MirAge provided high precision at the cost of a lower sensitivity.

Figure 1C shows the results of analyzing the HiSeq sample from Kraken with MirAge, together with the reported results from the Kraken and CLARK website, as well as the rerun analysis with both tools using the NCBI reference sequence database. To provide insight, three common threshold settings, used for the relative score of MirAge, are marked for both fast and sensitive modes.

### 4.1 Performance

For the 8 simulated sample sets, time measurements were taken on a machine with a 12 core Xeon Gold 6162 processor and 2TB RAM. Figure 1B visualizes the differences in run time in minutes between MirAge (both sensitive and fast mode), CLARK and Kraken2. The effects of the different algorithms reflect in performance, as each algorithm benefits differently from different types of content in datasets. MirAge proves to be competitive and performs well, favors towards precision.

With the current optimizations and due to the nature and size of the sample data, speed is limited to what performance hardware can deliver. By increasing memory or disk performance, a lower threshold becomes possible while maintaining a reasonable analyzing time, increasing overall sensitivity.

## 5 Discussion

We showed MirAge to be capable of consistently providing high precision taxonomic binning compared to the other tested tools and deliver comparable sensitivity on low complexity samples. Especially the high precision of MirAge enables users, such as in clinical environments, to benefit from the detailed results, as it allows to distinguish species in high detail. The ability to combine several reference databases enables versatile analyses while remaining capable of high throughput processing.

## Acknowledgements

We thank SurfSara for providing and maintaining their well-suited supercomputer environment and tailor-made support.

